# Demographic and genetic structure of a severely fragmented population of the endangered hog deer (*Axis porcinus*) in the Indo-Burma biodiversity hotspot

**DOI:** 10.1101/511154

**Authors:** Sangeeta Angom, Chongpi Tuboi, Mirza Ghazanfar Ullah, Syed Ainul Hussain

## Abstract

The population of the globally endangered hog deer (*Axis porcinus*) has declined severely across its geographic range. Intensive monitoring of its demographic and genetic status is necessary. Northeast India is a stronghold of the species; however, in recent years the population has been getting fragmented, and it is vulnerable to extinction. We examined the demographic and genetic structure of a small hog deer population in the floating meadows of Keibul Lamjao National Park (KLNP), located on the western fringe of the Indo-Burma biodiversity hotspot for conservation planning. We used a double-observer distance sampling method to derive the hog deer abundance and population structure. We also derived the genetic diversity of the population through microsatellite screening and bottleneck detection. Our study revealed that the abundance of the deer in the park was 1.82–4.32 individuals/km^2^. The adult male to female ratio and fawn to doe ratio were 36.2 ± 1.9 males/100 females and 16.5 ± 0.4 fawns/100 females, respectively. The mean number of alleles at 23 loci was 2.70 ± 0.18, the observed heterozygosity (H_O_) ranged from 0.26 to 0.63 (mean 0.42 ± 0.02), the expected heterozygosity (H_E_) ranged from 0.23 to 0.73 (χ = 0.51 ± 0.03), and the polymorphic information content (PIC) ranged from 0.2 to 0.67 (χ = 0.43 ± 0.03). The observed allelic distribution reveals that the population has not encountered any genetic bottleneck in the recent past. Although the population is declining, it still retains some rare alleles, and the genetic diversity is 50%. This diversity will probably not affect the short-term population growth but may affect the evolutionary potential by limiting the selection flexibility. Conservation measures coupled with a scientifically sound management regime may help the persistence of the population in the region at a time when the population still retains rare alleles and maintains reproductive fitness.

## Introduction

The hog deer (*Axis porcinus*) is a species endemic to South and Southeast Asia. There are two subspecies—the Southeast Asian subspecies (*A. p. annamiticus*) and the Indian subspecies (*A. p. porcinus*). The Southeast Asian subspecies originally occurred in China, Thailand, Laos, Cambodia and Vietnam and the Indian subspecies in Pakistan, Nepal, India, Bangladesh and Burma [1]. Thus it was once widespread, but the population has declined rapidly across its geographic range. It is estimated that the global decline rate over the last 21 years is 50% and that the species has declined by more than 90% within its Southeast Asian range [2]. The species has completely disappeared from Thailand, Laos and Vietnam [2].

In India, the species is distributed throughout the northern plains and in the Northeast [3]. The latter region has emerged as a stronghold for the species, with large local subpopulations in Assam and West Bengal [4] and in Manipur, where a small, secure subpopulation survives [5, 6]. Nevertheless, the species is estimated to have undergone a decline of 30–40% or more in the country over the last 21 years [7].

On the basis of these estimates (the average for the species in three generations exceeding 50%), the species is listed as “Endangered” in the IUCN Red List [7]. Hunting, habitat loss and habitat degradation have been the major drivers of the decline [2]. The extant population is, therefore, patchily distributed and highly fragmented [8]. The relict populations are found in low alluvial grasslands in the Brahmaputra flood plains and in Nepal, Bangladesh and Southwest Yunnan Province, in China [2]. These populations are probably suffering inbreeding depression [9, 10].

Genetic information from both mitochondrial and nuclear DNA suggests that the hog deer belongs to the subfamily Cervinae [11]. However, the phylogenetic position of *Axis porcinus* is controversial [12]. Classical taxonomy [13] and a phylogenetic study [11] place it in the genus *Axis*; however, other studies have pointed out that the species might be placed in the genus *Rusa* [12]. This issue may be resolved with more molecular and archaeological evidence [14]. Information on the genetic variability and structure is also critical for developing successful conservation strategies for this species. Little empirical information is available on the ecology and conservation genetics of this species.

Northeast India is particularly significant in terms of the lineage and genetic diversity of the hog deer, being a stronghold of the species [4] and possibly holding a link between the Indian subspecies and the Southeast Asian subspecies. The localised and isolated hog deer populations in the state of Manipur are not well studied, and, therefore, little is known about the status of these populations and their genetic structure. Recent studies have also revealed that the hog deer population in Keibul Lamjao National Park, Manipur is the Southeast Asian subspecies (*A. p. annamiticus*) [15]. Isolated populations such as these are considered more vulnerable to demographic, environmental and genetic stochasticity and therefore more vulnerable to local extinction [10, 16, 17, 18]. Understanding the population dynamics of any species requires a combination of demographic, genetic and ecological studies. The effective population size and the number of individuals that actually contribute to the gene pool are key factors in the dynamics [19]. In view of the foregoing, this study was carried out to examine the demographic status and genetic structure of the hog deer in Keibul Lamjao National Park (KLNP), Manipur, India.

## Materials and methods

### Study area

The study was conducted in KLNP, located on the southeastern fringes of Loktak Lake, between latitudes 24°27′N and 24°31′N and longitudes 93°53′E and 93°55′E, in the Barak-Chindwin river basin (Fig. 1). Loktak Lake is a wetland of international importance (Ramsar site) [20]. Presently the park has an extent of 40.05 km², of which 26.41 km² is covered by a thick and almost contiguous mat of floating meadows, which are locally known as *phumdis.* The remaining extent of 14.09 km² consists of open water, dry lands and hills [21]. The floating meadows of the park are of varying thickness, and they support small populations of the hog deer and Eld’s deer (*Rucervus eldii*). The construction of the Ithai barrage, in the early 1980s, has affected the natural process of meadow formation and led to rapid changes in the ecosystem of the lake [20]. The meadows used to settle down during the lean season and get replenished with soil and nourishment; now they float continuously, as a result of which their thickness has decreased and may not support the weight of the wild ungulates such as the hog deer and Eld’s deer, which are found sympatrically in the park.

**Fig. 1.** Map of Keibul Lamjao National Park, Manipur, India

The park experiences three distinct seasons, *viz.*, summer (March–May), the monsoon (June–October) and winter (November–February) [21]. The temperature ranges from a maximum of 34.4°C to a minimum of 1.7°C. The annual rainfall is 1460 mm. The humidity is highest in August, reaching up to 81% then, and least in March, when it is 49%. More than 185 species of herbs, grasses and sedges have been recorded from the meadows [22]. *Zizania latifolia*, *Phragmites karka*, *Saccharum munja*, *Hemarthria compressa*, *Leersia hexandra*, *Carex* spp., *Oryza rufipogon* and *Capillipedium* spp. are the major food plants of the wild ungulates of the park [6].

### Field methods

The entire park was divided into 1 km × 1 km grids to examine the distribution pattern and habitat use of the hog deer. Each grid was sufficiently large to encompass the range of a male or female hog deer. In each grid, two or three strip transects of length 500 m (n = 66 in 2006; n = 75 in 2007; n = 72 in 2008), were laid, according to the *phumdi* type and the presence of open water. Hog deer faecal pellet groups within 2 m of each transect were recorded. Habitat parameters were recorded in circular plots of radius 2 m along each transect at 50-m intervals: (1) ground cover (vegetation, litter, water and *phumdi*), (2) substratum character (water/*phumdi*/soil), (3) *phumdi* thickness, (4) water depth, (5) total number of faecal pellet groups, (6) signs of associated species, (7) ocular visibility, (8) distance from uplands/hills, (9) distance to water for drinking/wallowing and (10) disturbance factors (fishing, grazing and vegetable collection).

A 0.5 m × 0.5 m quadrat was laid within each plot to record the number of each plant species present. On the basis of the distribution of the faecal pellet groups, the different areas of the park were classified as high, medium and low-density areas.

Trial boat transect surveys were carried out at different study sites in the park to identify specific locations that were suitable for constructing *machans* (temporary bamboo watchtowers) for estimating and monitoring the hog deer population. This exercise provided baseline information for constructing *machans* and also helped classify the habitat into thick and thin *phumdis*. No *machan* was constructed in those blocks in which no faecal pellets were recorded. Twenty-two *machans* of height 7 m each were constructed.

The point count method [23, 24] was used to estimate the abundance of hog deer in a 20 km^2^ area of the park excluding the water bodies and thin *phumdis*. The high-pellet density areas in the park had eight *machans*, and the medium and low-pellet-density areas had seven *machans* each. Population monitoring study was conducted in the month of March from 2006 to 2008. The observations were made using 8 × 40 prismatic binoculars. Details such as number of individuals sighted, time of sighting, sighting distance, sighting angle, age, sex and habitat type were recorded.

To get more accurate estimates, the counts were replicated on five continuous days. Counting of animals was carried out when the grass had been cut and had dried and the visibility of the animals in the park was good. The animals were classified as adult male, sub-adult male, adult female, sub-adult female, juvenile and fawn [25, 26]. Field surveys were carried out between 0530 and 0900 hours and between 1500 and 1730 hours using a double-observer approach developed to estimate detection probabilities and to increase detection accuracy [27, 28]. During each point count, a designated “primary observer” indicates to the “‘secondary observer” all the animals detected, who then records all the detections. The observers take the primary and secondary roles alternately during the course of a survey. This approach permits estimation of observer-specific detection probabilities and abundance and is carried out under the assumption that no animal goes undetected during the sampling periods.

Another method, the drive count, was used to determine the accuracy of the point count technique. The drive count was carried out after the completion of the point count, at 0900 hours, when the animals were hidden in tall grass and thick bushes. During the drive counts, local people and forest department personnel were used to flush and count the animals at each point. The numbers obtained using the two methods were compared.

### Statistical analyses

The mean faecal pellet group density was calculated (number/km^2^, with the standard error) using MS Excel. Percentage area of the faecal pellet density (high, medium and low) was summarized. The area was then stratified according to the percentage area and faecal pellet distribution map was generated using the ArcView GIS software (9.1). The habitat data were pooled into a single matrix. Continuous independent variables were first evaluated for independence using Pearson correlations. Multiple regression models were run using SPSS 16 [29] to find the best combination of habitat variables that predicted hog deer habitat use as judged by explained variance, overall *F* and regression diagnostics. The data were normalised using the Z-transform. The scaling of the habitat use can be represented by a linear model of the form Y = a + bx_1_ + cx_2_ + dx_3_…, where Y is the predicted density and x represents the habitat variables.

The Distance 6.0 software package was used to derive the abundance [30]. The hog deer density was calculated using maximum peak sightings for 30 minutes to avoid duplicate counts. The best-fitting model on the basis of the Akaike information criterion (AIC) [31] was used to derive the density. The exponential rate of increase was derived by regressing the natural log (ln) of the population density (hog deer/km) against year. The exponential rate of increase (ṝ) is the slope of the regression line. The percentage change per year was calculated as percent change = (e^r^ − 1)*100 [32].

### Microsatellites genotyping

The total genomic DNA was extracted from tissue hair, antlers and faecal samples (n = 38) using Kit (QIAGEN), QIAamp® DNA Micro Kit (QIAGEN) and QIAGEN Stool Mini Kit, respectively, following the manufacturer’s protocol [33]. The DNA fragments were separated on an automated DNA sequencer (Applied Biosystems). Different individuals were genotyped as either homozygote or heterozygote on the basis of the band pattern shown for each microsatellite locus. GeneMapper v.3.7 software was used to analyse the allele frequencies, heterozygosity and other parameters of genetic variability. To control for allelic dropout (stochastic non-amplification of one allele), three or four PCR replicates were carried out for each sample according to the concentration of DNA in it [34]. Two repeats were performed, and if there was a heterozygote in three of four repeats, the site was considered to be a heterozygous locus; the same for a homozygote locus [35].

CERVUS 3.0 software was used to identify unique genotypes [36]. Samples that had mismatches at up to two loci were re-examined for possible genotyping errors or allelic dropout [37]. The probability of identity statistic, P(ID) [38], and a more conservative variant of P(ID), P(ID-sibs) [39] were calculated to ensure that the loci used could reliably discriminate related individuals.

CERVUS 3.0 software was used to derive the observed and effective number of alleles, allele diversity (observed and expected heterozygosity) and percentage of polymorphic loci [36]. The polymorphic information content (PIC) was calculated using the allelic frequencies. GENEPOP 4.7 software was used to derive the deviations from the Hardy-Weinberg equilibrium (HWE) [40].

### Bottleneck detection

Signatures of bottleneck events were investigated by comparing the expected heterozygosity for a sample (HE) with the heterozygosity that would be expected for a sample taken in a population at mutation-drift equilibrium with the same size and allele number (HEQ) [41]. As the allele number decreases faster than does the heterozygosity, a bottleneck is indicated by HE > HEQ in subsequent generations [42]. Transient heterozygosity excess and mode-shift tests was performed through 1000 simulations using the BOTTLENECK 1.2.02 software package [43], assuming that mutations of the microsatellite loci followed an infinite allele model (IAM), a stepwise mutation model (SMM) or a two-phase model (TPM). In the last case, it was assumed that 70% of the mutations consist of one step and 30% consist of multistep changes with a variance of 30 (default values). Tests were performed using three different methods (*viz*. the sign test, standardised differences test and Wilcoxon test) [42]. Once all the loci available in a sample were processed, the three statistical tests were performed for each mutation model [42, 44], and the allele frequency distribution was determined to see whether it was approximately L-shaped (as expected under a mutation-drift equilibrium) or not (recent bottlenecks provoke a mode shift).

## Results

### Distribution pattern

Faecal pellets were recorded in extents of 22.26, 22.04 and 22.73 km^2^ in the park during 2006-2008 respectively (Fig. 2). The high-density area (9.31 ± 0.30 km^2^) had 15.9 ± 1.0 pellets/km^2^, the medium-density area (8.10 ± 0.40 km^2^) had 6.66 ±1.30 pellets/km^2^ and the low-density area (4.8 ± 0.6 km^2^) had 2.61 ±0.15 pellets/km^2^. The overall pellet density was 0.34 ± 0.02 pellets/km^2^. The hog deer habitat usage was best explained by the combination of *phumdi* thickness, vegetation cover and abundance of short grass (R^2^ = 0.423; F = 11.475, df = 50, *p* < 0.001). The estimated coefficient of habitat use showed that the preference for a specific habitat by hog deer increased with increasing *phumdi* thickness (β = 0.087 ± 0.05), decreasing vegetation cover (β = −0.142 ± 0.044) and increasing abundance of short grass (β = 0.124 ± 0.048). In other words, it showed that hog deer favoured areas with thick *phumdi*, less vegetation cover and dominated by short grasses.

**Fig. 2.** Distribution pattern of hog deer in Keibul Lamjao National Park, India

### Detection probability

The model was selected once the data set was ready after truncation. Several robust models were considered on the basis of the AIC value. The criterion for model selection is that the detection function should satisfy [31] conditions [45] such as robustness of the function (shape, estimator efficiency). The probability of detection was estimated using eight models, *viz.*, the uniform cosine, uniform simple polynomial, half-normal cosine, half-normal simple polynomial, half-normal Hermite polynomial, hazard-rate cosine, hazard-rate Hermite polynomial and hazard-rate simple polynomial. The lowest AIC is an estimate of the best approximating model. Since the AIC value of the uniform cosine (*) was the lowest, it was chosen as the best model for the detection function to estimate the population density and abundance of the hog deer population (Table 1).

**Table 1.** Best model analysis of hog deer during population estimation (2006–08) in Keibul Lamjao National Park, India

| Model | AIC |  |  | D<sup>†</sup> |  |  | N<sup>†</sup> |  |  | P |  |  |
| --- | --- | --- | --- | --- | --- | --- | --- | --- | --- | --- | --- | --- |
|  | 2006 | 2007 | 2008 | 2006 | 2007 | 2008 | 2006 | 2007 | 2008 | 2006 | 2007 | 2008 |
| Uniform + simple polynomial | 344 | 305 | 319 | 3.01 | 2.33 | 2.17 | 57 | 47 | 43 | 0.54 | 0.72 | 0.69 |
| Uniform + cosine* | 94 | 72 | 75 | 2.94 | 2.75 | 2.51 | 65 | 61 | 57 | 0.91 | 0.97 | 0.95 |
| Half normal + cosine | 345 | 301 | 313 | 2.87 | 2.41 | 2.53 | 54 | 49 | 51 | 0.34 | 0.44 | 0.21 |
| Half normal + simple polynomial | 346 | 301 | 313 | 2.88 | 2.42 | 2.54 | 54 | 49 | 51 | 0.53 | 0.38 | 0.66 |
| Half- normal + Hermite polynomial | 345 | 301 | 313 | 2.87 | 2.41 | 2.53 | 54 | 49 | 51 | 0.57 | 0.70 | 0.48 |
| Hazard rate + cosine | 348 | 302 | 313 | 2.50 | 2.83 | 2.65 | 47 | 57 | 53 | 0.49 | 0.55 | 0.58 |
| Hazard rate + Hermite polynomial | 346 | 302 | 313 | 2.50 | 2.83 | 2.65 | 47 | 57 | 53 | 0.61 | 0.58 | 0.38 |
| Hazard rate + simple polynomial | 346 | 302 | 313 | 2.50 | 2.94 | 2.65 | 47 | 56 | 53 | 0.68 | 0.33 | 0.43 |

\*Uniform + cosine is the best fitted model based on the lowest AIC value. <sup>†</sup>D, density; N, number.

### Population trend

The encounter rates were 0.36 ± 0.33, 0.22 ± 0.20 and 0.24 ± 0.20 individuals/km^2^ during 2006-2008 respectively. However, the probabilities of detection were 0.38 ± 0.58, 0.31 ± 0.29 and 0.30 ± 0.25, and the effective detection radius (EDR) values were 1.92 ± 0.35, 1.19 ± 0.17 and 1.25 ± 0.17, respectively, during the three consecutive years. The mean cluster sizes were 1.53 ± 0.10, 2.29 ± 0.18 and 2.02 ± 0.18 during 2006-2008 respectively (Table 1). The estimated densities were 2.94 ± 0.57 (CV 19.5%), 2.75 ± 0.44 (CV 16.3%) and 2.51 ± 0.40 (CV 16.2%) individuals/km^2^ during 2006-2008 respectively, with a minimum of 1.82 individuals/km^2^ and a maximum of 4.32 individuals/km^2^ at 95% confidence level. The population size was 65 ± 12.6, 61 ± 9.9 and 57 ± 9.2 individuals, with a minimum of 44, 44 and 41 and a maximum of 96, 84 and 79 individuals at the 95% confidence level during 2006-2008 respectively. The maximum number of sightings was recorded between 0600 and 0630 hours at distances between 100-200m (Fig. 3 a & b). The population showed a declining trend from 2006-2008 (*p* < 0.05, R² = 0.916) (Fig. 3a).

**Fig. 3.** Frequency of sighting of hog deer during population estimation in Keibul Lamjao National Park, India: (a) frequency of sighting versus time and (b) frequency of sighting versus distance

The population structure was skewed towards adult females, followed by adult males (Fig. 4 a–c). The overall population structure also indicated a higher number of females compared with males, juveniles and fawns (Fig. 4d). The overall numbers of adult males, adult females and juveniles fluctuated, whereas the number of fawns declined steadily. The adult sex ratios were 34.2, 34.5 and 39.9 males/100 females and the doe to fawn ratios were 16.4, 17.2 and 15.8 fawns/100 females for the three consecutive years, respectively (Fig. 5). Overall, for the three study years, the male to female ratio was 36.2 ± 1.9 males/100 females, and the doe to fawn ratio was 16.5 ± 0.4 fawns/100 females respectively (Table 2).

**Fig. 4.** Population trend of hog deer in Keibul Lamjao National Park, India

**Fig. 5.** Age structure of hog deer population during population estimation in Keibul Lamjao National Park, India

**Table 2.** Density estimate of hog deer (2006–08) in Keibul Lamjao National Park, India

| Parameters | Estimates |  |  | % CV |  |  | 95% CI |  |  |  |  |  |
| --- | --- | --- | --- | --- | --- | --- | --- | --- | --- | --- | --- | --- |
|  |  |  |  |  |  |  | Minimum |  |  | Maximum |  |  |
|  | 2006 | 2007 | 2008 | 2006 | 2007 | 2008 | 2006 | 2007 | 2008 | 2006 | 2007 | 2008 |
| p* | 0.38 ± 0.58 | 0.31 ± 0.29 | 0.30 ± 0.25 | 15.6 | 9.59 | 8.48 | 0.27 | 0.25 | 0.25 | 0.52 | 0.37 | 0.38 |
| n/k* | 0.36 ± 0.33 | 0.22 ± 0.20 | 0.24 ± 0.20 | 9.33 | 10.3 | 10.5 | 0.29 | 0.18 | 0.18 | 0.44 | 0.27 | 0.28 |
| EDR* | 245 ± 19.2 | 240 ± 11.5 | 241 ± 10.2 | 7.83 | 4.8 | 4.24 | 208.9 | 217.8 | 220.8 | 287.8 | 265.6 | 263 |
| DS* | 1.92 ± 0.35 | 1.19 ± 0.17 | 1.25 ± 0.17 | 18.2 | 14.1 | 13.5 | 1.33 | 0.90 | 0.95 | 2.76 | 1.59 | 1.63 |
| ES* | 1.53 ± 0.10 | 2.29 ± 0.18 | 2.02 ± 0.18 | 6.74 | 8.2 | 8.87 | 1.33 | 1.93 | 1.68 | 1.75 | 2.72 | 2.42 |
| D* | 2.94 ± 0.57 | 2.75 ± 0.44 | 2.51 ± 0.40 | 19.46 | 16.3 | 16.2 | 1.99 | 1.99 | 1.82 | 4.32 | 3.80 | 3.46 |
| N* | 65 ± 12.6 | 61 ± 9.9 | 57 ± 9.2 | 19.46 | 16.3 | 16.2 | 44 | 44 | 41 | 96 | 84 | 79 |

\*p, probability of detection under the curve; n/k, encounter rate; EDR, effective detection radius; DS, group density; ES, group size; D, individual density; N, population.

### Genetic diversity in the wild

Of the 25 microsatellites screened, 23 were polymorphic, and two loci were non-amplified. The genetic variability study of the hog deer population (n = 27) showed that the number of alleles observed across the 23 loci varied from 2 to 5, whereas the mean number of alleles per locus was 2.70 ± 0.18. The average observed heterozygosity (Ho) was estimated at 0.42 ± 0.02, and the expected heterozygosity (He) was estimated at 0.51 ± 0.03. The mean PIC value was 0.43 ± 0.03. The heterozygosity is around 50%, which indicates a moderate genetic variation in the population (Table 3). Out of the 23 microsatellites, two loci, T193 and T507, deviated from the HWE (both *p*<0.001), and no deviation from the HWE was observed in the remaining 21 variable loci.

**Table 3.** Estimated numbers of hog deer in the various age and sex classes on the basis of the relative proportions seen during the population estimation exercise conducted in 2006–08 in Keibul Lamjao National Park, India and the estimated total population size

| Year | Adult male |  |  | Adult female |  |  | Juvenile |  |  | Fawn |  |  |
| --- | --- | --- | --- | --- | --- | --- | --- | --- | --- | --- | --- | --- |
|  | Mean | 95% lower CI | 95% upper CI | Mean | 95% lower CI | 95% upper CI | Mean | 95% lower CI | 95% upper CI | Mean | 95% lower CI | 95% upper CI |
| 2006 | 13 | 9 | 20 | 39 | 27 | 58 | 6 | 4 | 9 | 11 | 7 | 16 |
| 2007 | 12 | 9 | 17 | 35 | 26 | 49 | 2 | 2 | 3 | 6 | 4 | 8 |
| 2008 | 13 | 9 | 18 | 40 | 28 | 55 | 5 | 3 | 6 | 5 | 4 | 7 |
| Mean | 13 | 9 | 18 | 38 | 27 | 54 | 4 | 3 | 6 | 7 | 5 | 11 |
| SEM | 0.37 | 0.19 | 0.88 | 1.34 | 0.86 | 2.69 | 1.01 | 0.66 | 1.55 | 1.67 | 1.05 | 2.64 |

### Bottleneck detection

Of the 23 loci found to be polymorphic in the wild population, only seven loci showed a significant heterozygosity excess (Table 4). The three statistical tests were performed for each mutation model [42, 44]. Overall, the Wilcoxon test showed no evidence of a bottleneck in the past (*p =* 0.894 for IAM; *p =* 0.993 for TPM; *p =* 0.999 for SMM) (Table 5). The global analysis indicated no significant heterozygosity excess (H_*E*_<H_*EQ*_): 0.252 ± 0.025 < 0.284 ± 0.018 (under IAM); 0.252 ± 0.025 < 0.323 ± 0.021 (under TPM); 0.252 ± 0.025 < 0.359 ± 0.026 (under SMM). The hog deer population did not encounter a genetic bottleneck in the recent past as the allele frequency distribution had an approximately L-shaped distribution (as expected under a mutation–drift equilibrium). Although the hog deer population has gradually declined in recent years, the population has retained some of its rare alleles and a 50% genetic diversity in the wild.

**Table 4.** Details of microsatellite loci tested on wild population of *Axis porcinus* at Keibul Lamjao National Park, India (a total of 27 individual samples were used in this study)

| Name of locus | Repeat type | Size range | T<sub>a</sub><sup>†</sup> | N<sub>A</sub><sup>†</sup> | H<sub>O</sub>/H<sub>E</sub><sup>†</sup> | HWE/LD<sup>†</sup> | PIC<sup>†</sup> | PID<sub>ID(Sibs)</sub><sup>†</sup> | Reference |
| --- | --- | --- | --- | --- | --- | --- | --- | --- | --- |
| Ca18* | Di | 194–212 | 52 | 5 | 0.63/0.73 | Yes/No | 0.675 | 0.421 | Gaur et al. 2003 |
| T507* | Tetra | 174–186 | 58 | 3 | 0.44/0.65 | Yes/No | 0.568 | 0.479 | Jones et al. 2002 |
| ILSTS005* | Di | 180–182 | 50 | 2 | 0.577/0.49 | Yes/No | 0.366 | 0.605 | Zhang et al. 2005 |
| BM4208* | Di | 162–164 | 54 | 2 | 0.370/0.475 | Yes/No | 0.358 | 0.615 | Say et al 2005 |
| T123* | Tetra | 148–152 | 58 | 2 | 0.259/0.23 | Yes/No | 0.2 | 0.793 | Jones et al. 2002 |
| T108* | Tetra | 128–154 | 58 | 2 | 0.556/0.51 | Yes/No | 0.375 | 0.593 | Jones et al. 2002 |
| Ca42* | Di | 164–178 | 60 | 2 | 0.407/0.44 | Yes/No | 0.338 | 0.638 | Gaur et al. 2003 |
| OarFCB193* | Di | 100–126 | 54 | 2 | 0.26/0.33 | Yes/No | 0.272 | 0.715 | DeYoung et al 2003 |
| Cervid1* | Di | 144–168 | 54 | 4 | 0.56/0.71 | Yes/No | 0.642 | 0.437 | DeYoung et al 2003 |
| BM6506* | Di | 198–200 | 54 | 2 | 0.37/0.45 | Yes/No | 0.346 | 0.629 | Anderson et al 2002 |
| RT27* | Di | 130–144 | 56 | 2 | 0.33/0.41 | Yes/No | 0.321 | 0.659 | Poetsch et al. 2001 |
| CelJP27 | Di | 154–164 | 59 | 4 | 0.519/0.72 | Yes/No | 0.647 | 0.434 | Coulson et al 1998 |
| MAF70 | Di | 124–128 | 50 | 3 | 0.44/0.64 | Yes/No | 0.559 | 0.486 | Zhang et al. 2005 |
| RT6 | Di | 98–104 | 50 | 3 | 0.407/0.6 | Yes/No | 0.526 | 0.512 | Poetsch et al. 2001 |
| BM4107 | Di | 152–190 | 50 | 3 | 0.33/0.51 | Yes/No | 0.449 | 0.575 | Zhang et al. 2005 |
| T193 | Tetra | 176–186 | 59 | 3 | 0.407/0.53 | Yes/No | 0.443 | 0.566 | Jones et al. 2002 |
| RT1 | Di | 206–208 | 54 | 2 | 0.26/0.23 | Yes/No | 0.2 | 0.793 | Poetsch et al. 2001 |
| T156 | Tetra | 128–130 | 59 | 2 | 0.35/0.45 | Yes/No | 0.343 | 0.632 | Jones et al. 2002 |
| CSSM16 | Di | 164–166 | 55 | 2 | 0.407/0.33 | Yes/No | 0.272 | 0.715 | Zhang et al. 2005 |
| INRA011 | Di | 212–220 | 54 | 3 | 0.444/0.56 | Yes/No | 0.449 | 0.55 | DeYoung et al 2003 |
| D-F/R | Di | 154–190 | 54 | 3 | 0.5/0.68 | Yes/No | 0.588 | 0.466 | Anderson et al 2002 |
| INRABERN185 | Di | 102–114 | 52 | 4 | 0.41/0.66 | Yes/No | 0.675 | 0.473 | Zhang et al. 2005 |
| NVHRT48 | Di | 102–106 | 52 | 2 | 0.37/0.43 | Yes/No | 0.33 | 0.648 | Poetsch et al. 2001 |
|  |  |  |  | **2.70 ± 0.18** | **0.42 ± 0.02/0.51 ± 0.03** |  | **0.43 ± 0.03** | **0.58 ± 0.02** |  |

\*Loci required for unambiguous individual identification.
<sup>†</sup>Ho, observed heterozygosity; He, expected heterozygosity; PIC, polymorphic information content; HWE, Hardy-Weinberg equilibrium; PID, probability of identity, the probability that two different individuals will share the same multilocus genotype at a given number of loci; PID<sub>ID(Sibs)</sub>, the probability that a pair of siblings will share the same genotype; Ta, annealing temperature; N<sub>A</sub>, number of alleles; LD, linkage disequilibrium.

**Table 5.** Bottleneck detection of hog deer in Keibul Lamjao National Park, India indicating (H_*E*_<H_*EQ*_) mutation–drift equilibrium obtained with three mutation models

| Loci | Observed | IAM* |  | TPM* |  | SMM* |  |
| --- | --- | --- | --- | --- | --- | --- | --- |
|  | He* | Heq* | Prob* | Heq* | Prob* | Heq* | Prob* |
| AY302223 | 0.207 | 0.362 | 0.27 | 0.421 | 0.14 | 0.484 | 0.055 |
| BM4208 | 0.331 | 0.211 | 0.298 | 0.24 | 0.358 | 0.265 | 0.402 |
| RT6 | 0.268 | 0.353 | 0.383 | 0.427 | 0.2 | 0.488 | 0.088 |
| INRA011 | 0.234 | 0.368 | 0.295 | 0.406 | 0.182 | 0.488 | 0.063 |
| RT1 | 0.14 | 0.22 | 0.481 | 0.231 | 0.427 | 0.264 | 0.345 |
| Ca42 | 0.171 | 0.215 | 0.511 | 0.238 | 0.461 | 0.243 | 0.45 |
| Cervid1 | 0.326 | 0.48 | 0.212 | 0.542 | 0.085 | 0.606 | 0.024 |
| RT27 | 0.171 | 0.214 | 0.514 | 0.242 | 0.466 | 0.249 | 0.432 |
| NVHRT48 | 0.171 | 0.22 | 0.531 | 0.237 | 0.474 | 0.26 | 0.405 |
| BM6506 | 0.073 | 0.225 | 0.33 | 0.251 | 0.262 | 0.251 | 0.235 |
| OarFCB193 | 0.257 | 0.22 | 0.395 | 0.239 | 0.445 | 0.249 | 0.48 |
| T123 | 0.23 | 0.205 | 0.397 | 0.234 | 0.486 | 0.249 | 0.518 |
| T156 | 0.382 | 0.221 | 0.246 | 0.243 | 0.286 | 0.261 | 0.338 |
| CelJP27 | 0.205 | 0.361 | 0.255 | 0.419 | 0.136 | 0.482 | 0.046 |
| D-F/R | 0.336 | 0.366 | 0.447 | 0.423 | 0.319 | 0.485 | 0.157 |
| T193 | 0.297 | 0.357 | 0.414 | 0.426 | 0.235 | 0.481 | 0.127 |
| T108 | 0.107 | 0.202 | 0.476 | 0.241 | 0.354 | 0.257 | 0.309 |
| T507 | 0.532 | 0.361 | 0.207 | 0.423 | 0.281 | 0.476 | 0.396 |
| BM4107 | 0.208 | 0.372 | 0.275 | 0.424 | 0.136 | 0.485 | 0.047 |
| MAF70 | 0.507 | 0.361 | 0.284 | 0.42 | 0.402 | 0.478 | 0.514 |
| L23481 | 0.292 | 0.211 | 0.335 | 0.235 | 0.402 | 0.262 | 0.468 |
| AF232760 | 0.307 | 0.208 | 0.311 | 0.239 | 0.376 | 0.253 | 0.422 |
| INRABERN185 | 0.037 | 0.212 | 0.232 | 0.235 | 0.175 | 0.252 | 0.17 |
|  | 0.252 ±<br>0.025 | 0.284 ±<br>0.018 |  | 0.323 ±<br>0.021 |  | 0.359 ±<br>0.026 |  |

\*He, expected heterozygosity; Heq, heterozygosity equilibrium; Prob, probability; IAM, infinite allele model; TPM, two-phase model; SMM, stepwise mutation model.

## Discussion

KLNP lies in the Indo-Burma biodiversity hotspot, one of the most biologically rich regions of the world, with only 5% of the natural habitat of the hotspot remaining [46, 47]. The Indo-Burma hotspot is also one of the most densely populated regions. Thus it has experienced rapid economic development and changing consumption patterns, which has put the natural ecosystems of the hotspot under immense anthropogenic pressure. KLNP is a very small area within this hotspot that is under tremendous anthropogenic pressure. Consequently its biodiversity and habitats are exposed to various forms of risk. The hog deer population in the park is facing conservation issues such as hunting, poaching, predation and competition with domestic livestock. Furthermore, the population is threatened by the periodic loss of habitat due to flooding and development projects, lack of connectivity with other viable habitats, a high risk of site-level extinction.

The hog deer is typically shy, nocturnal in habit [48] and highly sensitive to human disturbances. Thus it is absent in areas where there are dense human populations and in agricultural areas [49]. The declines in its population are attributed to large-scale transformations in the native range, mainly due to agricultural developments [1]. Considering the anthropogenic pressures in KLNP, the long-term viability of this small population of hog deer seems uncertain. Moreover, KLNP, consisting of an extent of only 40 km^2^ dominated by floating meadows and open waters [6, 22], appears to be insufficient to support any growth in the population. This necessitates conservation and management efforts towards securing a viable population of hog deer in KLNP.

The threats to the species in KLNP are further intensified by its small population size [19], the highly isolated nature of the park and the lack of connectivity between KLNP and the surrounding habitats. The present study found a declining trend in the hog deer population of KLNP, from 186 individuals in 2003 [50] to 57 individuals in 2008, with densities estimated at 2.93, 2.75 and 2.51 individuals/km² with upper confidence intervals of 4.32, 3.80 and 3.46 individuals/km² over the three sampling years, respectively. This declining population trend may be a consequence of low and declining numbers of fawns, with the fawn to female ratios being 16.4, 17.2 and 15.8 fawns/100 females over the sampling period. The reduced numbers of fawns may be due to predation by stray dogs from nearby settlements.

The consequences of habitat fragmentation for the genetic structure and variability are not yet well understood [51]. However, several studies have pointed out that fragmented and isolated populations such as the one of the hog deer in KLNP may be susceptible to increased population differentiation due to loss of heterozygosity, genetic drift and inbreeding depression [9, 10, 52, 53]. A 50% loss of genetic diversity was observed in the population, with indications that some of its rare alleles have been retained. Factors other than genetic diversity are also imperative for the persistence of a species that has become reduced in numbers. These factors include demographic and environmental stochasticity and natural catastrophes [54]. With respect to the hog deer population in KLNP, unless inbreeding depression is a factor [55], genetic diversity will probably not affect the short-term population growth but may affect the evolutionary potential by limiting selection flexibility [56].

Genetic diversity helps populations adapt to changing environments. With higher levels of variation, it is more likely that some individuals in a population will possess variations of alleles that are adaptive to the changing environment. The population will persist for more generations because of the success of these individuals [57]. In our study, the *H*_*E*_ of the hog deer population was 0.51 ± 0.03, close to the mean *H*_*E*_ of 0.559 of sika deer in China but much higher than the mean *H*_*E*_ of the Sichuan sika deer (0.477). It was slightly lower than the mean *H*_*E*_ of Przewalski’s gazelle (*Procapra przewalskii*) (*H*_*E*_ = 0.552) [58], Manchurian sika deer (0.584) and South China sika deer (Jiangxi, 0.585; Zhejiang, 0.589) [59]. Compared with other ungulates, such as the red deer (*Cervus elaphus*) (*H*_*E*_ = 0.78 [60]; *H*_*E*_ = 0.804 [61]; *H*_*E*_ = 0.62–0.85 [62]), forest musk deer (*Moschus berezovskii*) (*H*_*E*_ = 0.8–0.9) [63] and Mongolian wild ass (*Equus hemionus hemionus*) (*H*_*E*_ = 0.83) [64], the hog deer had a relatively low *H*_*E*_ value and was equivalent to the milu (*Elaphurus davidianus*) (*H*_*E*_ = 0.46–0.54) [65]. The PIC is a measure of the variation between microsatellite loci. It is generally considered that a PIC value greater than 0.5 represents highly polymorphic loci, a PIC value less than 0.5 and greater than 0.25 represents moderately polymorphic loci, and a PIC value less than 0.25 represents low polymorphic loci [66]. In our study, the PIC values of eight of the 23 microsatellite loci were higher than 0.5, indicating highly polymorphic loci, while 13 were moderately polymorphic loci and two were low polymorphic locus. The heterozygosity and PIC indicated that the hog deer in KLNP had moderate genetic diversity (50%) (Table 4).

Hartl and Clark [67] considered that population deviations from the Hardy-Weinberg equilibrium (HWE) are mainly due to small populations, not random mating, gene mutation, migration or other factors. In the present study, two loci out of 23 microsatellites, T193 and T507, deviated significantly from the HWE (both *p*<0.001). No deviation from the HWE was observed in the remaining 21 variable loci. This can be attributed to the reduction in the population size of hog deer in the area. In addition, due to the relatively closed geographical position of KLNP, *i.e.*, the lack of connectivity to the surrounding areas, very few hog deer were distributed in its periphery [68]. This indicated that the moderate genetic variation came from inside the sampling area due to geographical barriers and anthropogenic factors.

Microsatellite polymorphism studies of sika deer in China found that 16 microsatellite loci significantly deviated from the HWE, but the *H*_E_ values were higher than the *H*_O_ values [59]. In the present study, the *H*_O_ values were significantly higher than the *H*_E_ values. When a population experiences a reduction in its effective size, it generally develops a heterozygosity excess at selectively neutral loci [42]. Xu et al. [69] used microsatellites to detect the population structure of Himalayan marmots and found a heterozygosity excess due to a historic population decline. Wu et al. [59] tested bottlenecks in four Chinese sika deer species and found that the Sichuan sika deer had not experienced a population bottleneck in recent history.

The study also revealed that the hog deer population of KLNP has not encountered any genetic bottleneck in the recent past. The bottleneck investigation through a gene diversity excess test showed that the expected heterozygosity under equilibrium was higher than the heterozygosity observed (Table 5). The global analysis indicated no significant heterozygosity excess (H_*E*_<H_*EQ*_): 0.252 ± 0.025 < 0.284 ± 0.018 (under IAM); 0.252 ± 0.025 < 0.323 ± 0.021 (under TPM); 0.252 ± 0.025 < 0.359 ± 0.026 (under SMM) (Table 5). The null hypothesis of a mutation–drift equilibrium was rejected, which suggests that there is no apparent genetic signature of a population bottleneck in the recent past. The mode shift test also resulted in a normal L-shaped allele distribution curve, indicating the presence of a larger proportion of alleles in the low allele frequency classes and thus a lack of recent genetic bottlenecking in the hog deer population at KLNP.

This study assessed the population structure and genetic diversity of the small and isolated population of hog deer in the Barak-Chindwin basin, and it highlighted the need to prioritize conservation measures for the persistence of the population in its natural habitat. Although the hog deer population is showing a steadily declining trend over the years, the genetic variability of the population was moderate, which indicates that there is a high probability of the population persisting if remedial measures are undertaken. However, if effective conservation measures coupled with scientifically sound management regimes are not adopted for this population, further reduction of the genetic variation in the future cannot be ruled out.

## Acknowledgements

This study was conducted under the project “Conservation Ecology of Eld’s Deer and Its Wetland Habitat”, sponsored by the Wildlife Institute of India. We are grateful to the Department of Forests, Government of Manipur for granting us logistic support. We thank the Director and the Dean at the Wildlife Institute of India for their support. We thank Dr. Ruchi Badola for all the discussions and critical input at different stages of the study.

## Author contributions

**Conceptualization:** Syed Ainul Hussain

**Data curation:** Sangeeta Angom, Chongpi Tuboi

**Formal analysis:** Sangeeta Angom

**Funding acquisition:** Syed Ainul Hussain

**Investigation:** Sangeeta Angom, Syed Ainul Hussain

**Methodology:** Sangeeta Angom,Syed Ainul Hussain

**Project administration:** Syed Ainul Hussain

**Resources:** Syed Ainul Hussain

**Supervision:**Syed Ainul Hussain

**Validation:** Syed Ainul Hussain

**Writing—original draft:** Sangeeta Angom, Chongpi Tuboi

**Writing—review and editing:** Syed Ainul Hussain

